# Advancing Pediatric Sleep Assessment through Spectral Sleep Scoring: A Comparison with Polysomnography

**DOI:** 10.64898/2026.08.28.747921

**Authors:** Alessandra E. Shuster, Julie A. Onton, Neal Nakra, Katharine C. Simon

## Abstract

Understanding how sleep changes across development requires methods that are both efficient and sensitive to maturational differences in sleep physiology. Spectral sleep scoring is a promising approach, but has primarily been validated in adult populations. This study presents and validates a spectral scoring approach for pediatric sleep. Participants (n = 192; 9-18y) completed overnight in-lab sleep studies. Visual scoring was determined by registered polysomnographic technologists and confirmed by board certified sleep physicians following AASM guidelines. Visual stages included: wake, non-rapid eye movement (NREM) stage 1 (N1), NREM stage 2 (N2), NREM stage 3 (N3), and rapid-eye movement (REM) sleep. Spectral activity within each epoch was assessed for the following spectral stages: Wake (40–95 Hz), Light (~11– 15.5 Hz), Hi Deep (1–3 Hz), Lo Deep (0.1–1 Hz), and REM (~17–26 Hz). Concordance rates were checked across agreement matrices between visual and spectral stages, with expected high agreement between: 1) visual Wake and spectral Wake; 2) visual N2 and spectral Light; 3) visual N3 and spectral HiDeep/LoDeep; and 4) visual REM and spectral REM. Cohen’s κ was calculated from concordance matrices. There was high agreement between visual and spectral stages (71-89%, κ=.76). Our results provide evidence that spectral scoring is a promising method for efficiently scoring pediatric sleep. Moreover, this method provides additional, clinically relevant information that is not readily captured through traditional visual scoring. This may be particularly valuable for assessing sleep physiology in pediatric clinical populations, where developmental differences in EEG patterns may complicate visual scoring.

## Introduction

Sleep during childhood and adolescence is characterized by ongoing neurodevelopmental maturation, resulting in systematic differences in both sleep macrostructure and the underlying electroencephalographic (EEG) spectral features.^1,2^ Thus, it is critical that approaches to characterize pediatric sleep account for these dynamic age-related changes in sleep physiology. Current American Academy of Sleep Medicine (AASM) guidelines for sleep scoring acknowledge age-dependent differences, specifying pediatric scoring criteria distinctly from adult scoring.^3,4^ AASM guidelines remain the gold standard for sleep scoring, however, the visual scoring process is time-intensive, with an overnight 8-hour PSG taking up to two hours to score by qualified professionals.^5^ An estimated 855,000 sleep disorder diagnoses occur per year in U.S. youth ages 0-18 years.^6^ While not all sleep disorders require a PSG study for diagnosis, the high prevalence of disorders, coupled with potential for repeat PSG studies or PSG studies not resulting in diagnosis, likely represent a substantial scoring burden. Thus, there is an urgent need for efficient methods to evaluate pediatric sleep and support large-scale clinical and research applications.

Early efforts to automate sleep scoring were motivated by the desire to improve efficiency as well as address the challenges associated with visual scoring in clinical contexts, where pathological features can complicate the accurate interpretation of sleep recordings.^7,8^ To date, numerous sleep scoring algorithms have been developed, however the majority have been designed and validated primarily in adult populations.^5,9^ Consequently, sleep scoring algorithms developed and validated in adult populations may not generalize well to pediatric sleep, given the dynamic maturational changes observed across time. Indeed, sleep undergoes substantial changes across late childhood and adolescence (~9-18 years), mirroring pubertal and cortical maturation.^1,2^ During this time, sleep shifts towards shorter duration, less time in non-rapid eye movement (NREM) stage 3 (N3), and a proportional increase in NREM stage 2 (N2).^1,10^

Sleep macrostructure changes are accompanied by developmental shifts in microstructure. Sleep spindles (12-15 Hz) increase in frequency, but decline in amplitude which parallels declines in slow wave activity (SWA; delta <4Hz) amplitude associated with cortical changes persisting through late adolescence.^2,11,12^ Spindles and SWA, which predominate in frontal regions during late childhood, progressively shift towards posterior regions across adolescence.^1,13^ Both are key EEG features used in scoring N2 and N3 sleep, which together comprise approximately three quarters of the night during adolescence.^14^ REM sleep also shows adolescent spectral shifts, with greater power in younger ages for all frequencies below 15 Hz excepting the alpha range,^1^ which may further complicated generalization of adult-trained algorithms to pediatric populations.

A limited number of automated sleep scoring algorithms have been developed to assess pediatric electroencephalography (EEG); however, several key limitations remain. One pediatric algorithm demonstrated good agreement for overall sleep-wake distinction, but poor reliability for individual sleep stages.^15^ Other algorithms are promising, but often rely on multiple channels and struggle with more complex EEG.^16–18^ Beyond these limitations, existing algorithms are designed primarily to reproduce conventional visual scoring. While visual staging provides critical characterization of sleep macrostructure, it represents a coarse summary of underlying EEG. Additionally, AASM visual guidelines rely on categorical pediatric vs. adult criteria, switching to adult scoring at age 18, whereas sleep undergoes continuous maturation rather than an abrupt developmental shift across childhood and adolescence. Sleep microstructure also contains developmentally and clinically-relevant spectral distinctions not captured in visual staging alone. For instance, visual scoring does not distinguish between delta (<4 Hz) activity and slow oscillations (SOs; <1Hz) in the scoring of N3. Yet, prior research suggests that SOs and delta represent distinct brain activity,^19^ have separable roles in memory,^20^ and differ in their spatial organization, with SOs marked by larger amplitude global waves and delta by smaller amplitude local waves.^20–23^ SOs are also considered a direct marker of brain maturation and connectivity during childhood^24^, are associated with memory outcomes,^25^ and demonstrate alterations between healthy and clinical pediatric populations.^26,27^ Together, these findings demonstrate that pediatric sleep EEG contains developmentally meaningful spectral information beyond what is captured in visual staging.

Despite existing advances, automated sleep scoring algorithms remain uncommon in the analysis of routinely collected clinical pediatric PSG. This limited adoption underscores the need for continued development and validation of pediatric-specific sleep scoring algorithms, as well as collaboration between clinicians and researchers to facilitate implementation of these tools at scale. The present study aimed to modify and validate an automated spectral sleep scoring algorithm, previously validated in adult PSG,^28^ in pediatric PSG. This algorithm characterizes sleep by quantifying dominant spectral activity, generating epoch-level sleep scores that distinguish subtle patterns of EEG activity that may be difficult to detect by visual inspection of raw EEG, upon which conventional sleep scoring relies. By distinguishing epochs dominated by <1 Hz activity separately from 1-3 Hz activity, it preserves neurophysiological information that is pertinent to developmental brain maturation. In modifying this algorithm for pediatric populations, our goal is to provide an efficient supplementary tool to clinical sleep scoring, which provides developmentally-relevant information beyond traditional scoring.

## Materials and Methods

### Participants

This study included a retrospective sample of 192 youth (9-18 years, Table 1, Figure 1) who underwent clinically-indicated sleep studies at Rady Children’s Health, Orange (RCHO) between 1/1/2018-9/30/2025. This sample size was determined based on an a prior power analysis conducted in G*Power for overall agreement between visual and spectral scoring. Based on findings in adults indicating overall agreement of approximately 70% with this spectral algorithm,^28^ a sample size of 178 was determined sufficient to detect agreement exceeding a minimum threshold of 60% at α = 0.05 and 80% power. The final sample size of 192 was achieved through retrospective chart review, with additional records included beyond the minimum required sample to account for potential exclusions due to short PSG recordings (<4 hours). Participants were generally healthy, with no major medical or neurodevelopmental comorbidities, severe psychiatric disorders (e.g. schizophrenia, bipolar, etc.), sleep disorders with severe symptoms (e.g. severe obstructive sleep apnea (OSA), severe insomnia, narcolepsy), neurological conditions, or history of significant brain injury or loss of consciousness. Patients with autism spectrum disorder and/or attention-deficit/hyperactivity (ADHD) disorder, or who were taking medications known to impact sleep were also excluded. Patients with mild symptoms in the following categories were eligible for inclusion: sleep disordered breathing, mild-to-moderate OSA, asthma, anxiety, depression, obesity, hypertension, allergic rhinitis, and tonsil/adenoid conditions. This retrospective study and procedures were approved by the Institutional Review Board at Rady Children’s Health, Orange with reliance at University of California, Irvine.

**Table 1:** Demographics Table. Table showing demographic information across participants.

| Demographic | Metrics |
| --- | --- |
| Age | 13.36 ± 2.69 years (see Figure 2) |
| Sex | F = 90<br>M = 102 |
| Race/Ethnicity | White/Caucasian: 101<br>Asian: 14<br>Black/African American: 2<br>Multiracial: 9<br>Other: 53<br>Not reported: 13 |

**Figure 1.**
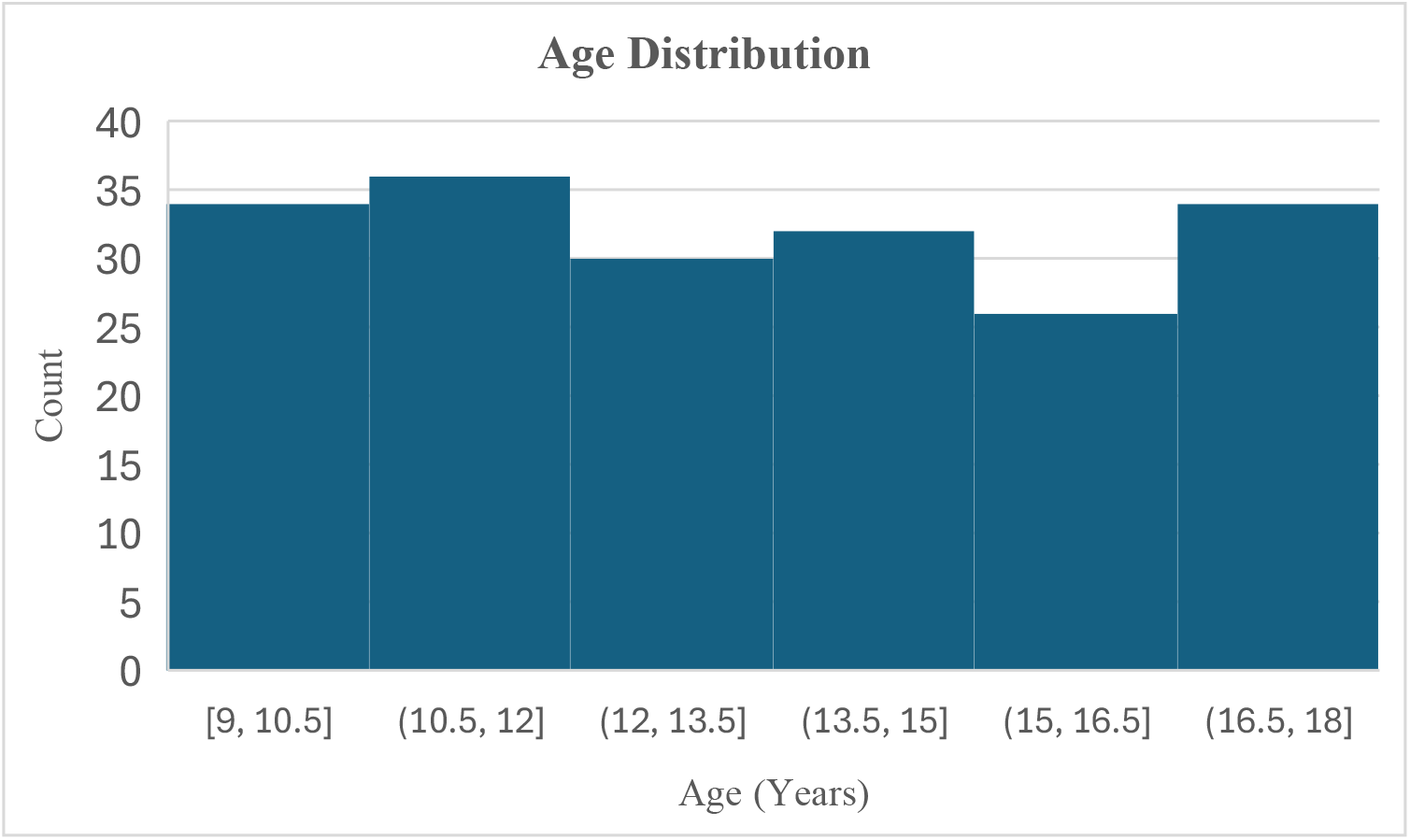
Distribution of participant ages (years).

### Polysomnography

All participants completed a clinically-indicated outpatient sleep study with overnight polysomnography monitoring including 6 neural channels (F3, F4,C3, C4, O1, O2) recorded via Nihon Kohden PSG-1100 amplifier at 200 Hz sampling rate. On-line filtering during recording included an analog 3rd-order Butterworth filter and a software 2nd-order Butterworth filter. Post-acquisition, data was processed in Polysmith and neural channels were re-referenced to the contralateral mastoid. Visual sleep scoring was determined by registered polysomnographic technologists and confirmed by a board certified sleep physician following AASM guidelines ^4^. EEG recordings were then exported from Polysmith as EDFs and further processed in Matlab using EEGLAB (2022.1). In cases where recordings commenced before lights-off or continued beyond lights-on, EDF files were trimmed to exclude periods during which room lights were on.

### Spectral scoring

Following the methods of Onton and colleagues (2016, 2024),^28,29^ the algorithm uses Morlet wavelets to decompose EEG data per neural channel from 0.1-90 Hz. The number of wavelets increases smoothly from 3 cycles to 30 cycles from lowest to highest frequencies to optimize tradeoff between temporal and frequency resolution. Power values were assessed in decibels (dB) as follows: 10∗ log10 (amplitude), baseline-corrected by averaging across the entire night excluding high-noise epochs, and used to create a baseline-normalized spectrogram (for more details see: Onton et al., 2016, 2024).^28,29^ Spectral scores were assigned per 30 second epoch by evaluating power across predefined frequency ranges. Spectral stages were categorized into Wake (40–85 Hz), Light (10–15.5 Hz), HiDeep (1–3 Hz), LoDeep (0.1-1Hz), and REM (17– 25 Hz), which is expected to roughly correspond to visually scored Wake, NREM 2, NREM 3 (both HiDeep and LoDeep), and REM sleep respectively, though spectral scores were not created to necessarily align with visual scoring (Table 2). Per epoch, the conditional probability of the participant being in each spectral stage is calculated, and the stage with maximal probability is assigned as the spectral score. While the algorithm was run for all neural channels, analyses focused on frontal channels (F3 and F4) given that the original spectral algorithm was optimized on frontal electrodes and that sleeping brain rhythms reflecting developmental brain maturation, such as spindles and SWA, predominate in frontal regions.^1,13^

**Table 2:** Spectral stages (column 1) and their associated spectral frequency ranges (column 2) alongside the associated target visual stages (column 3).

| **Spectral Stage** | **Spectral Frequency** | **Target Visual Stage** |
| --- | --- | --- |
| Wake | 40-85 Hz | Wake |
| Light | 10-15.5 Hz | NREM 2 |
| HiDeep | 1-3 Hz | NREM 3 |
| LoDeep | 0.1-1 Hz | NREM 3/SOs* |
| REM | 17-25 Hz | REM |

\*Slow Oscillation Events

The frequency ranges above incorporated two modifications to the original adult-validated algorithm to better reflect pediatric sleep physiology. First, target spindle range was defined at 10-15.5 Hz, lowered from 11-15.5 Hz to account for slower spindle frequencies observed in younger populations.^12^ Second, REM spectral frequency range was set at 17-24 Hz, adjusted from 16-30 Hz, based on literature showing a dominance of low beta (15-24 Hz) vs. high beta (25-30 Hz) in adolescents.^30^ The lower boundary was set at 17 Hz to avoid overlap with spindle range and based on preliminary observations that no adult REM beta peaks fell below 17 Hz.^29^ Within these bounds, individualized REM frequency was determined per record, based on peak frequency ± 2 Hz following the original algorithm. Similarly, individualized spindle frequency was also determined based on peak frequency within the target spindle range ± 1.5 Hz.

Two additional updates were made to adapt the algorithm for pediatric populations. The first was based on the standard progression of overnight sleep stages in healthy children and adolescents, where REM sleep follows NREM within a sleep cycle.^31^ Initial optimization checks of the algorithm indicated that it incorrectly scored spectral REM during instances scored as visual Wake at the beginning of some records. Thus, the algorithm was updated to correct spectral REM to Wake in instances where REM had been spectrally scored before the first epochs of NREM. The second update was made to account for high-amplitude K-complexes during visually scored N2 sleep, which produce prominent <1 Hz activity.^32,33^ Given that K-complex amplitude is greater in younger populations,^33^ this dominant <1 Hz activity sometimes results in spectral classification as LoDeep. While LoDeep is indeed intended to capture <1 Hz activity, inconsistent K-complex activity within an epoch should not rise to the level of LoDeep, but in populations with sparse true LoDeep, even modest increases in <1 Hz activity can be scored as LoDeep. Thus, an update was made to screen for consistent slow waves (low variability) compared to solo K-complex spikes (high variability). Specifically, spectral LoDeep epochs were reclassified as Light sleep when variability in <1 Hz amplitude exceeded an individualized threshold. For each participant, this threshold was determined by assessing average <1 Hz amplitude variability across all epochs, with LoDeep epochs exceeding 1 standard deviation above the mean variability reclassified as Light sleep. This threshold was determined by initial optimization analyses indicating that this threshold yielded best agreement with visual scores.

### Comparison of visual and spectral scoring

Visual and spectral scores were compared by evaluating epoch-by-epoch scores for each and assessing agreement (see Figure 2 for example visual comparison). Agreement was determined between all stages, populating a confusion matrix between the five visual stages (Wake, N1, N2, N3, REM) and spectral stages (Wake, Light, Hi Deep, Lo Deep, REM).

**Figure 2.**
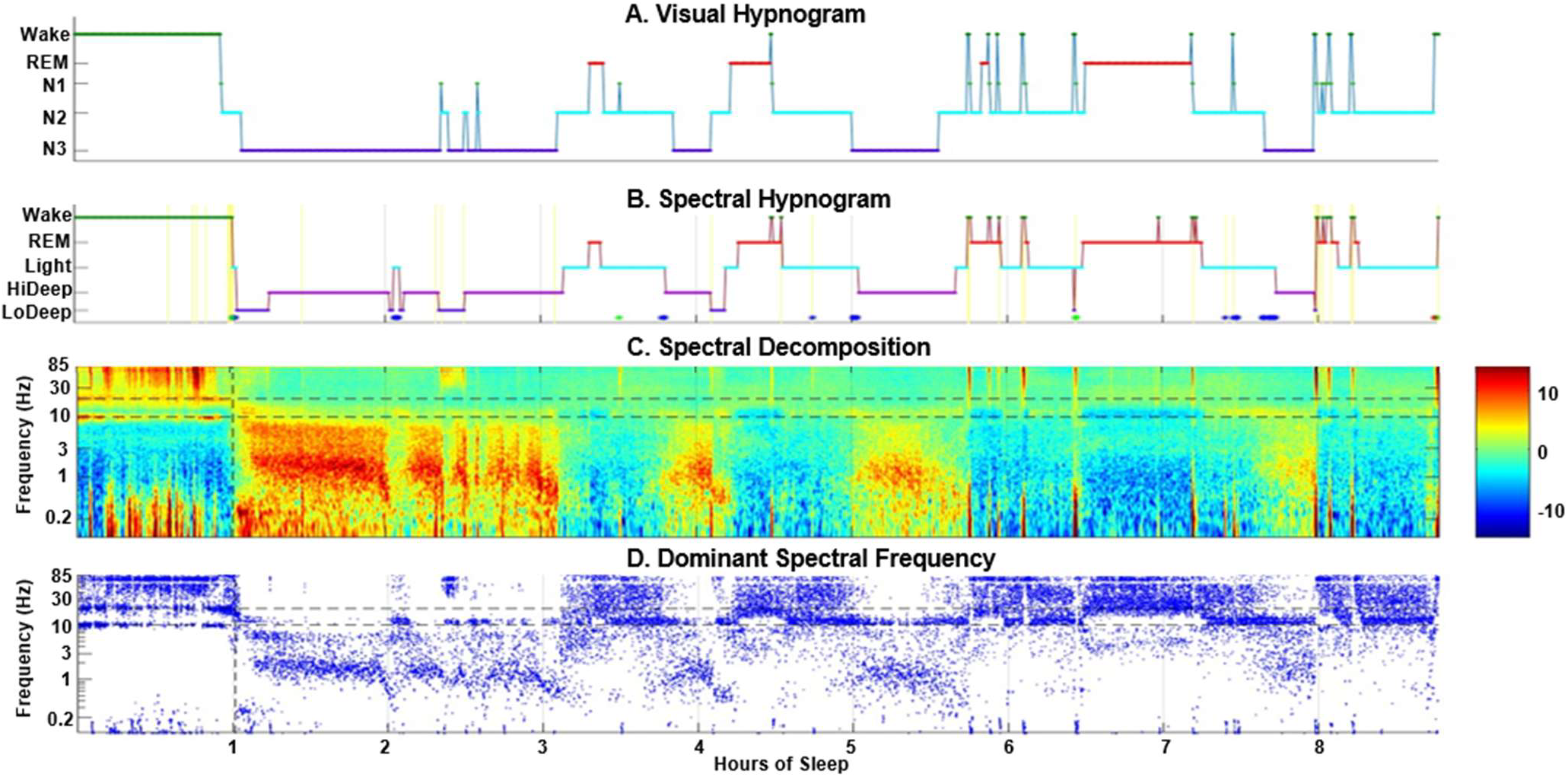
Example report showing alignment between visually scored hypnogram (A) and spectrally scored hypnogram (B) for one record. C: Depicts spectral decomposition of power (legend) across time (x-axis) by frequency (y-axis). D: Depicts dominant spectral frequency (y-axis) across time (x-axis).

Agreement was expected to align between: 1) Visual Wake and Spectral Wake; 2) Visual N2 and Spectral Light; 3) Visual N3 and Spectral HiDeep; 5) Visual REM and Spectral REM (Table 2). As spectral LoDeep captures <1 Hz activity matching SO events, it was expected that this spectral stage would match visual N3 as well. Confusion matrices were calculated at the individual subject level as counts, and then summed across all subjects to create an overall agreement matrix. Cohen’s *κ* was calculated from the matrix to assess overall agreement. To further understand how the algorithm compared to visual scoring, the agreement counts were converted to overall percentages using two complementary normalizations. First, matrices were normalized by visual stage such that each cell was divided by the total number of epochs in the corresponding visual category. For example, agreement between visual N2 and spectral Light sleep was defined as the number of epochs scored as both visual N2 and spectral Light, divided by the total number of epochs scored as visual N2. These will be referred to as visual-normalized values going forward. Second, matrices were also normalized by spectral stage, using the same approach, but with each cell divided by total number of epochs in the corresponding spectral category. These will be referred to as spectral-normalized values going forward.

## Results

Across all records, 296,863 epochs were assessed representing almost 2500 hours of sleep. Participants spent similar proportions of the night in visual and spectral Wake (16% vs. 17%), visual N2 and spectral Light sleep (44% vs. 36%), visual N3 and spectral HiDeep (19% vs. 21%), and visual and spectral REM (16% vs. 20%), respectively (Table 3). Figure 2 provides a representative example of the alignment between visual scoring (Figure 2a) and spectral scoring (Figure 2b), with corresponding spectral activity and dominant frequency across the night (Figure 2c-d). Agreement counts between all visual and spectral stage pairings are reported in Table 4. The distribution of sleep stages was broadly similar between visual and spectral scoring (Table 3). With the exception of spectral LoDeep, normalized confusion matrices reflected high percent agreement across all expected visual-spectral pairings (~71-89%) under both visual- and spectral-normalized conditions (Table 5a-b).

**Table 3:**
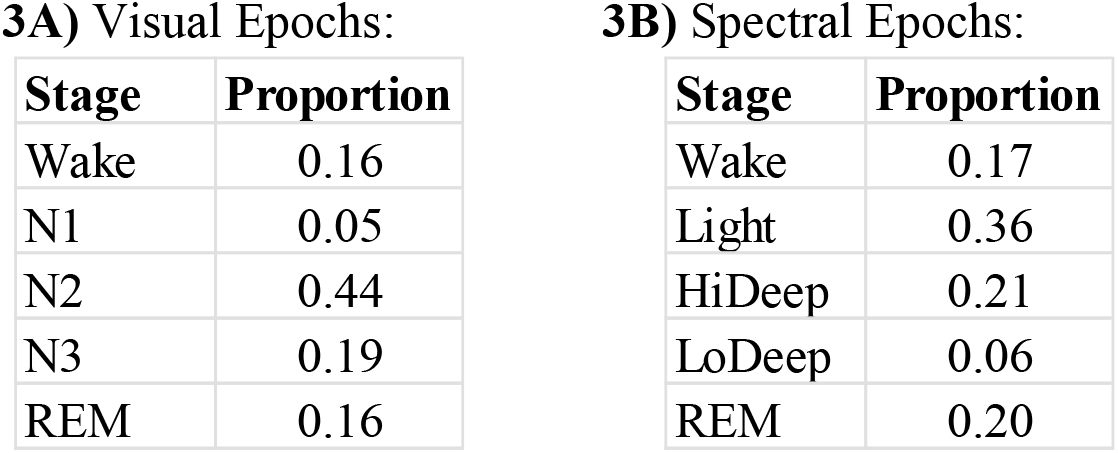
Proportion Stages Scored Across All Epochs: Proportion of time spent in each visual sleep stage (2a) and each spectral sleep stage (2b) across the night.

**Table 4:** Agreement Counts. Concordance matrix showing counts (epochs) of visual scores (rows) and spectral scores (columns).

|  | Wake | Light | HiDeep | LoDeep | REM |
| --- | --- | --- | --- | --- | --- |
| Wake | 36283 | 4294 | 774 | 1425 | 3626 |
| N1 | 5301 | 4614 | 501 | 959 | 3824 |
| N2 | 7221 | 92663 | 12535 | 10354 | 7670 |
| N3 | 179 | 3214 | 47913 | 5221 | 10 |
| REM | 1629 | 3279 | 127 | 189 | 43058 |

**Table 5:**
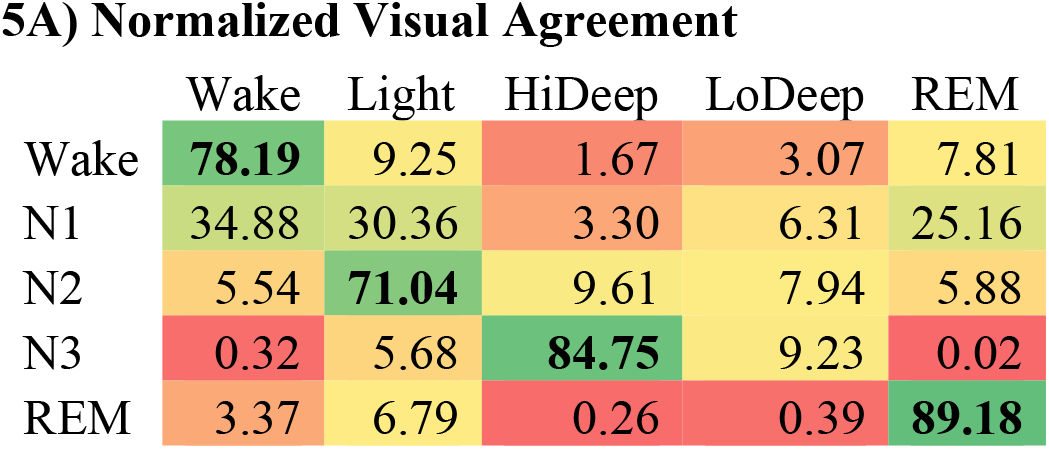
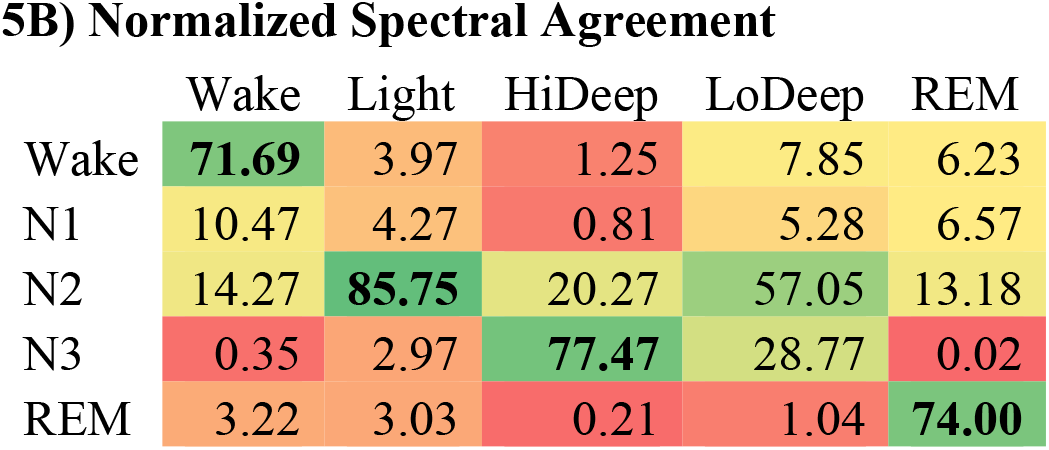
Agreement Percentages. a: Concordance matrix showing percent agreement between visual and spectral scores, normalized by visual scores. b: Concordance matrix showing percent agreement between visual and spectral scores, normalized by spectral scores.

Highest concordance was found for REM sleep in the visual-normalized matrix, where spectral REM correctly identified visual REM at a rate of 89% (Table 5a). In the spectral-normalized matrix, visual REM accounted for 74% of spectral REM (Table 5b). Demonstrating second highest concordance, spectral Light appropriately signified visual N2 86% of the time (Table 5b). Conversely, under visual-normalization, visual N2 matched spectral Light 71% of the time (Table 5a). High concordance was also found for visual N3 and spectral HiDeep, at 85% under visual-normalization (Table 4a) and 77% under spectral-normalization (Table 5b). LoDeep was not scored frequently across records (6%, Table 5b) and did not have high agreement with any visual stage (1-57%, Table 5a-b).

Due to the absence of a spectral equivalent for visual N1 and the relatively low proportion of epochs scored as spectral LoDeep (6%, Table 5b), Cohen’s kappa was calculated across four visual stages (Wake, N2, N3, REM) and four corresponding spectral stages (Wake, Light, HiDeep, REM). The resulting Cohen’s κ indicated agreement at κ = .76 (95% CI = .761 - .765). This level of agreement can be interpreted as substantial, consistent with established benchmarks for Cohen’s κ ranges of 0.61–0.80 indicating strong concordance between visual and spectral scoring beyond chance.^34^

## Discussion

The present study offers a promising approach for efficiently evaluating pediatric sleep, that holds potential for large-scale clinical and research applications where rapid sleep assessment is essential. Our pediatric-adapted spectral algorithm demonstrated substantial agreement with visual scoring (κ = 0.76), with strong concordance ranging between 71-89% across visual-spectral stages: 1) Wake-Wake; 2) N2-Light; 3) N3-HiDeep; 4) REM-REM. Notably, the pediatric algorithm demonstrated improved agreement relative to the original adult validation across all of these pairings.^28^ This may be due to a more robust sample size in the present study and tailored pediatric modifications. Interestingly, whereas the adult validation found that visual N3 was accounted for by spectral HiDeep and LoDeep combined (81%),^28^ the present study found that visually scored N3 in youth was captured reliably by HiDeep alone (85%). This is likely due to relatively few epochs (6%) scored as LoDeep in the present study, which may reflect the often observed ‘first night effect’ associated with undergoing a sleep study in a new environment, resulting in disruption to the depth and quality of sleep.^35^ An alternative explanation could be that yet uncharacterized sleep alterations are present in this pediatric cohort with mild medical conditions. Slow-wave sleep fragmentation and delta power alterations have been documented in children with respiratory conditions, although these findings are typically observed in greater disease severity.^36,37^ Thus, this spectral scoring approach may elucidate more subtle alterations in sleep associated with mild clinical pathology and should be pursued in future studies.

A distinct advantage of this spectral algorithm is that it characterizes dominant neural dynamics underlying each epoch, preserving important spectral distinctions not captured by conventional scoring. This approach may be particularly valuable in pediatric clinical settings, where pathology can disrupt neurodevelopment and sleep physiology. For instance, youth with Duchenne and Becker muscular dystrophy (DMD) exhibit an age-related decline in slow oscillation density compared with typically developing controls, whereas spindle activity is not altered.^38,39^ By distinguishing between <1 Hz slow oscillatory and 1-4 Hz delta activity, this algorithm provides an efficient tool for quantifying what is growingly recognized as distinct physiological processes,^22^ with potential to improve understanding of neurodevelopment in rare conditions such as DMD. More broadly, this approach also holds promise for common pediatric-onset conditions. For instance, in attention deficit-hyperactivity disorder (ADHD), findings are mixed regarding alterations to sleep macrostructure, whereas alterations to SWA are consistently reported, with reduced frontal SOs implicated in compromised executive function.^40,41^ Together, these findings highlight the potential of this algorithm as an efficient tool to provide clinically meaningful spectral characterizations overlooked by conventional staging. Ultimately, this spectral staging approach could be integrated into hospital-based sleep studies, providing rapid summaries of underlying sleep physiology as a supplement to the more time-intensive visual scoring.

Beyond hospital applications with traditional polysomnography, this spectral approach holds relevance for remote EEG assessment. A growing number of wearable EEG devices have increased accessibility to capture sleep physiology in ecologically valid home environments, also making longitudinal assessments more feasible.^42^ This is particularly relevant for pediatric populations, where full understanding of developmental changes in both healthy and clinical populations requires long-term within-subject assessment.^10^ Given the frontal predominance of sleeping brain rhythms associated with neurodevelopmental maturation,^1,13^ integration of this algorithm with frontal forehead EEG devices can enable widespread and scalable assessment of pediatric sleep physiology. As the field advances, such integrations can facilitate efficient longitudinal monitoring of pediatric sleep and facilitate new insights into developmental trajectories outside of hospital and traditional laboratory settings.

### Limitations

Participants were relatively healthy neurotypical youth referred for clinical polysomnography. While some had mild medical conditions, including mild sleep-disordered breathing (see Methods for full inclusion), this did not appear to compromise spectral sleep staging, demonstrated by the high agreement between visual and spectral scores in this sample. However, it is possible that the relatively few epochs of LoDeep (6%) observed in the current cohort reflects the presence of mild medical pathology. Nevertheless, these findings suggest that spectral features underlying sleep staging remain reliable in this subset of pediatric clinical populations, highlighting this algorithm as a feasible approach for clinical PSG assessment. It is important to note that this validation may not generalize to patients with neurological disorders, significant sleep pathology, or other major medical comorbidities. For instance, this algorithm was developed in consideration of typical overnight sleep architecture in youth, where NREM sleep precedes the first instance of REM.^31^ Thus, it would require re-evaluation for disorders such as narcolepsy where REM may pathologically occur before NREM. Similarly, additional evaluation is needed in patient populations with impacted EEG activity, such as seizure disorders where epileptiform activity presents challenges for visual scoring. Future refinement of the algorithm in more varied pediatric populations will further improve its clinical applications.

## Conclusion

Collectively, the present validation demonstrates the potential of this spectral sleep scoring algorithm as a complementary approach to conventional staging. By integrating developmentally informed spectral measures with clinical relevance, this framework holds potential for large-scale use across hospital, laboratory, and home-based settings. This approach represents an important step toward more efficient characterization of pediatric sleep across development in healthy and clinical populations.

## Data Availability

Data is available upon reasonable request.

## Declaration of generative AI and AI-assisted technologies in the manuscript preparation process

No AI was used in the preparation of this work.

## Acknowledge

We thank Lia Galut, Kalysa Bui, and Estella Knobbe for their assistance in data collection.

